# RL-118 and 11β-HSD1 target engagement through TAPS assay: behaviour and molecular analysis

**DOI:** 10.1101/2020.10.27.356881

**Authors:** D. Puigoriol-Illamola, J. Companys-Alemany, N. Homer, R. Leiva, S. Vázquez, D. Mole, C. Griñán-Ferré, M. Pallàs

**Affiliations:** Department of Pharmacology, Toxicology and Therapeutic Chemistry. Pharmacology Section. Faculty of Pharmacy and Food Sciences. University of Barcelona, Av Joan XXIII 27-31, 08028 Barcelona, Spain; Institute of Neuroscience, University of Barcelona (NeuroUB), Passeig de la Vall d’Hebron 171, Barcelona, Spain; Mass Spectrometry Core, Edinburgh Clinical Research Facility, Queen’s Medical Research Institute, Edinburgh, United Kingdom; Medicinal Chemistry Section. Department of Pharmacology, Toxicology and Therapeutic Chemistry. Faculty of Pharmacy and Food Sciences, University of Barcelona, Av. Joan XXIII, 27-31, 08028 Barcelona, Spain; MRC Centre for Inflammation Research, Queen’s Medical Research Institute, University of Edinburgh, Edinburgh, United Kingdom

## Abstract

Taking into consideration the convergence of ageing, stress and neurodegenerative diseases, such as AD, there is impaired GC signalling. Therefore, the study of GC-mediated stress response to chronic moderate stressful situations, as account in daily life, becomes of huge interest to design pharmacological strategies to prevent neurodegeneration.

To address this issue, SAMP8 were exposed for 4 weeks to the CMS paradigm and treated with RL-118, an 11β-HSD1 inhibitor. In fact, several pieces of evidence link the inhibition of this enzyme with reduction of GC levels and cognitive improvement, while CMS exposure has been associated with reduced cognitive performance. The aim of this project was to assess whether RL-118 treatment could restore the deleterious effects of CMS on cognition and behavioural abilities, but also on molecular mechanisms that compromise healthy ageing in SAMP8 mice.

On the one hand, we determined the target engagement between RL-118 and 11β-HSD1. Therefore all the beneficial effects previously described in SAMP8 treated with the drug can undoubtedly be attributed to the inhibition of this enzyme. Besides, herein we observed decreased DNA methylation, hydroxymethylation and histone phosphorylation induced by CMS but, on the contrary, increased after RL-118 treatment. In addition, CMS exposure produced ROS damage accumulation, and increments of pro-oxidant enzymes as well as pro-inflammatory mediators through NF-κB pathway and astrogliosis markers, like *Gfap. Of* note, those modifications were recovered by 11β-HSD1 inhibition. Remarkably, although CMS altered mTORC1 signalling, autophagy was increased in SAMP8 treated with RL-118 mice. Also, we found amyloidogenic APP processing pathway favoured and decreased synaptic plasticity and neuronal remodelling markers in mice under CMS, but changed after RL-118 treatment. In consequence, detrimental effects on behaviour and cognitive performance were detected in CMS exposed mice, but restored after concomitant 11β-HSD1 inhibition by RL-118.

Overall, CMS is a feasible intervention to understand the influence of stress on epigenetic mechanisms underlying cognition and accelerating senescence. However and most important, 11β-HSD1 inhibition through RL-118 turned up to restore the majority of these detrimental effects caused by CMS, indicating that GC excess attenuation may become a potential therapeutic strategy for age-related cognitive decline and AD.

## 2. Introduction

In the past few years, stress and stress response have been an overwhelming public health issue with a huge amount of literature, and in turn misinformation of the detrimental effects stress may have in modifying aging and in the development of several diseases. It is highly supported that stress is a key determinant of healthy and pathological aging of the brain (Catania et al., 2009; Sotiropoulous et al., 2008). Drug discovery studies have become of paramount importance in order to offer different drugs to treat or prevent the detrimental effects on cognition during aging. In consequence, diverse tools have recently been developed considering the drug-target binding, as well as cellular permeability, specificity and cytotoxicity, such as the toxicity-affinity-permeability-selectivity (TAPS) assay that is based on overexpressing the protein of interest and treating the cells with the drug, followed by FACS (fluorescence-activated cell sorting) coupled to mass spectrometry (MS) analysis.

Stressful situations activate a neuroendocrine response, which leads to the release of catecholamines at the first stage, and later with glucocorticoids (GCs). Active GCs bind to their receptors promoting slow genomic actions as well as rapid nongenomic effects, such as glucose release, lipolysis, motivation to eat palatable food and up-regulation of the expression of anti-inflammatory cytokines (Pazirandeh et al., 2002; Sandi, 2013). The ability to mount GC release is determined by the quality, intensity and chronicity of the stressful stimulus (Dhabhar, 2018). Primarily, stressful experiences are adaptive, facilitating the restoration of physiological and behavioural homeostasis and necessary for the establishment of enduring memories. However, when stressful situations are persistent in time, memory formation and reasoning become impaired (Tatomir et al., 2014; Wang et al., 2016). In this context it has been suggested that the deleterious effects of stress and GC are transient in nature, because stress-induced hippocampal atrophy and hippocampal-dependent behaviour may be reversed after a stress-free period. Overall, it is proposed that stress and GC can have widespread detrimental effects on mood and cognition, by inducing changes in brain structure (involving the generation and loss of neurons and dendritic atrophy) and function (electrophysiological activity and cellular signalling) (Sotiropoulous et al., 2011).

Several studies suggest that environmental stressors influence hypothalamic-pituitary-adrenal (HPA) axis activity and behaviour by altering the methylation status of key genes concerned with the regulation of the stress response (Harman & Martín, 2019; Sotiropoulous et al., 2008). Recent evidence has demonstrated significant associations between epigenetic alterations and stress showing that under chronic stressors influence, histone acetylation and DNA methylation are decreased, among others (Harman & Martín, 2019; Puigoriol-Illamola et al., 2020a).

Moreover, prolonged exposure to GCs has been related to immunosuppression, metabolic syndrome, diabetes, osteoporosis, reproductive failure, hypertension and mood and affective disorders. Mal-adaptive adjustments to stress may, sequentially, lead to symptoms of depression and Alzheimer’s disease (AD). Additionally, recent clinical studies suggest that GCs are implicated in the pathogenesis and/or progression of AD (Sotiropoulous et al., 2011). In particular, chronic stress has been associated with increased Aβ deposits and hyperphosphorylated Tau (Bisht et al., 2018; Puigoriol-Illamola et al.,2020a). In view of these affections, it is important to note that stressful events can have long-term consequences for the immune system. On one hand, pro-inflammatory cytokines have been implicated in the genesis of AD, becoming part of the amyloid plaques and triggering Aβ production. Additionally, NF-κB appears to be essential as it triggers a feed-forward cycle in which increased cytokine levels result in resistance to the GC immunosuppression, leading to further increases in cytokine release and, therefore, activation of the HPA axis (Irwin & Miller, 2007; Pace et al., 2007). Increasing evidence has reported a link between the HPA axis and Oxidative Stress (OS) in reactive oxygen species (ROS) production (Bonet-Costa et al., 2016; Schiavone et al., 2013), another factor related to AD pathology and progression.

In addition, GCs are also involved in the regulation of cell fate, modulating pro-/anti-apoptotic mechanisms and survival proteins, such as BDNF, CREB, Bcl2 and NCAM, so that may exert significant influence over neuroplasticity (Pittenger & Duman, 2008; Sandi, 2004, et al., 2013; Sotiropoulous et al., 2008; Wang et al., 2016). Noteworthy for cell survival and longevity, autophagy activation becomes crucial, as it participates in the elimination of disrupted proteins and is implied in the maintenance of cellular homeostasis (Puigoriol-Illamola et al., 2020a).

Disposition of active GCs is controlled by 11β-hydroxysteroid dehydrogenase type 1 (11β-HSD1) enzyme and up to now, our group has demonstrated that its inhibition leads to neuroprotective effects (Leiva et al., 2017; Puigoriol-Illamola et al., 2018, 2020b). The aim of this project is to assess RL-118 - 11β-HSD1 target engagement and evaluate whether 11β-HSD1 inhibition is able to face the detrimental effects of CMS.

## 3. Material and Methods

### Cloning

The E2-Crimson-human HSD11B1 gene (variant 1) was synthesised by GenScript in vector pUC57. The DNA sequence for E2-Crimson was sourced from Clontech. The gene for the fluorescent-HSD11B1 was ligated into vector pcDNA3.1 (Invitrogen) using restriction sites NheI (N-terminus) and NotI (C-terminus) (Fig. S1).

### Transient expression of fluorescent target protein

HEK293 cells were passaged in poly-D-Lysine treated plates and incubated overnight in OPTI-MEM medium (Lonza) at 37°C, 5% CO2. The cells were transiently transfected the following day with pcDNA3.1-E1-Crimson-huHSD11B1 DNA using Lipofectamine 2000 (Invitrogen) in OPTI-MEM medium by standard transfection protocol. The transfected cells were maintained at 37°C, 5% CO_2_ for 24 h post-transfection before the TAPS assay was performed. Transfection of the cells and compound screening were performed in a 6-well plate format.

### TAPS Assay

#### Compound incubation

RL-118 drug was diluted to a concentration of 20 μM (DMSO<1%) in tissue culture medium (DMEM with 10% FBS, 1% L-Glutamine, 1% penicillin-streptomycin, (Life Technologies)), then incubated with transfected cells for three hours at 37°C, 5% CO_2_. 20 μM compound concentration was selected as an appropriate concentration for assay development and screening. Following incubation, the cells were detached from the plate by gentle pipetting and centrifuged for 5 min at 1000 rpm. Culture medium was removed by pipetting and the cell pellet re-suspended in FACS buffer (PBS + 2% FBS) to wash off unbound compound. The cells were centrifuged as above, and the wash buffer removed. The cells were then re-suspended in FACS buffer and transferred to 5 ml FACS tubes. The tubes were wrapped in foil and placed on ice until sorting.

### FACS sorting

Cells were sorted using BD FACS Aria II system fitted with 100 μm nozzle. Data was acquired and processed using BD FACS Diva Software version 8.0.1. Cell fluorescence was detected using the 640 nm laser for E2-Crimson excitation. The filter used was 670/14 nm detecting fluorescence emission of E2-Crimson in the 663-677 nm range. Cells were sorted and collected into four cell populations defined by fluorescence intensity, forward (FSC) and side scatter (SSC) gating was used to exclude dead cells and only live cells were collected. Cells were collected in 5 ml FACS tubes containing 1 ml of DMEM with 10% FBS, 1% L-Glutamine and 1% penicillin-streptomycin. Each population of cells was centrifuged at 1000 rpm for 5 min. Medium was removed and the cell pellet resuspended in 20 mM HEPES pH7.0. The suspension was briefly sonicated to lyse the cells before centrifugation, as before, to pellet cell debris. The lysate (supernatant) was transferred to LC-MS vials and stored at −20°C prior to MS analysis.

#### Mass Spectrometry detection of the compounds in cell lysates

The chromatographic and mass spectrometer used was the SCIEX Triple Quad 5500+ LC-MS/MS System-QTRAP (Triple Quad^™^). 10μl injection of the cell lysate was loaded directly onto a HSST3 (150 x 2.1 mm, Thermo Fisher Scientific) column at a high flow rate, causing the proteinaceous material to flow to waste. A series of valve switches led to the elution of the extracted sample from the column directly onto the analytical column. Solvent A was water with 0.1% formic acid and solvent B was methanol with 0.1% of formic acid. Automated tune settings were used to achieve the maximum ion signal for the analyte for initial validation experiments, optimising on tube lens voltage, parent to product transitions and collision energy for transition. Peaks detected in this initial scan were then identified by molecular weight and checked versus their mass-charge ratio. Data was acquired and processed using Sciex OS-MQ software.

### Animals

Female SAMP8 mice (n=48) 4 months old were used to carry out behavioural, cognitive and molecular analyses. We divided these animals into four groups: SAMP8 Control (Control, n=12), SAMP8 treated with RL-118 (11β-HSD1i, n=12), SAMP8 under CMS (CMS, n=12) and SAMP8 treated with RL-118 under CMS (SAMP8 11β-HSD1i+CMS, n=12). Animals had free access to food and water and were kept under standard temperature conditions (22±2°C) and 12h: 12h light-dark cycles (300lux/0 lux). RL-118 was administered at 21 mg/kg/day by oral gavage for 4 weeks. Chronic Mild Stress (CMS) consisted of several different stressful stimulus applied to the corresponding animals daily during 4 weeks.

Studies and procedures involving mice brain dissection and subcellular fractionation were performed following the institutional guidelines for the care and use of laboratory animals established by the Ethical Committee for Animal Experimentation at the University of Barcelona.

### Chronic Mild Stress Treatment

CMS procedure used in the present study has been previously validated in SAMR1 and SAMP8 mice (Puigoriol-Illamola et al., 2020b). For 4 weeks, mice were daily exposed to various randomly scheduled, low-intensity environmental stressors. The number and the type of stressful stimuli applied changed everyday as well as the sequence of the stressors in order to guarantee the degree of unpredictability (Puigoriol-Illamola et al., 2020a). Stressful stimuli included 2 h of physical restraint, 24 h of sawdust removal, 24 h of food deprivation, 24 h of water deprivation, 24 h of wet bedding, 1 min of tail nipping at 1 cm from the tip of the tail and overnight illumination.

### Behavioural and cognitive tests

#### Elevated Plus Maze (EPM)

The Elevated Plus Maze (EPM) was performed as previously described (Griñán-Ferré et al., 2016). It is based on mice preference for dark enclosed places, therefore it evaluates anxiety-related and risk-taking behaviours. EPM apparatus consists of 2 opened and 2 closed arms elevated 50 cm from the floor. Mice were placed on the central platform, facing opened arms and allowed to explore the apparatus for 5 min. After that, mice were returned to their home cages and EPM arms were cleaned with 70% ethanol in order to avoid any olfactory clues. Behaviour was scored with SMART^®^ ver. 3.0 software and each trial were recorded for later analysis. Parameters evaluated included time spent on opened and closed arms, rearing, defecation and urination.

#### Novel Object Recognition Test (NORT)

The Novel Object Recognition Test (NORT) protocol employed was as described in Puigoriol-Illamola et al. (2020b). In brief, mice were placed in a 90°, two-arms, 25-cm-long, 20-cm-high, 5-cm-wide black maze. Before performing the test, the mice were individually habituated to the apparatus for 10 min for 3 days. On day 4, the animals were submitted to a 10-min acquisition trial (first trial), during which they were placed in the maze in the presence of two identical, novel objects at the end of each arm. After a delay (2h and 24h), the animal was exposed to two objects one old object and one novel object. The Time that mice explored the Novel object (TN) and Time that mice explored the Old object (TO) were measured. A Discrimination Index (DI) was defined as (TN-TO)/(TN+TO). To avoid object preference biases, objects were counterbalanced. The maze, the surface, and the objects were cleaned with 70% ethanol between the animals’ trials to eliminate olfactory cues.

#### Morris Water Maze (MWM)

This test evaluates both learning and spatial memory (Vorhees, 2006). An open circular pool (100 cm in diameter, 50 cm in height) filled with water was used. Water was painted white with latex in order to make it opaque and its temperature was 22± 1°C. Two main perpendicular axes were established (North-South and East-West), thus configuring four equal quadrants (NE, NW, SE, and SW). Four visual clues (N, S, E, W) were placed on the walls of the tank so that the animal could orientate and could fulfil the objective. The test consists of training a mouse to find a submerged platform (Learning phase) and assesses whether the animal has learned and remembered where the platform was the day that it is removed (Test). The training lasts five consecutive days and every day five trials are performed, which have a different starting point (NE, E, SE, S, and SW), with the aim that the animal recognizes the visual clues and learns how to locate the platform, avoiding learning the same path. At each trial, the mouse was placed gently into the water, facing the wall of the pool, allowed to swim for 60 seconds and there was not a resting time between trials. If the animal was not able to locate the platform, the investigator guided it to the platform and was allowed to rest and orientate for 30 seconds. The platform was placed approximately in the middle of one of the quadrants, 1.5 cm below the water level. Above the pool there was a camera that recorded the animals’ swimming paths and the data was analysed with the statistical program SMART^®^ ver.3.0. During the learning phase, a learning curve was drawn, in which is represented the latency to find the platform each training day. On the day test, more parameters were measured, such as the target crossings and the swum distance in the platform zone.

### Brain Processing

Three days after the behavioural and cognitive tests, 12 animals per group were euthanized for protein extraction, RNA and DNA isolation. Brains were immediately removed from the skull and the hippocampus was isolated, frozen on powdered dry ice and maintained at −80°C until procedures.

### Western Blotting

Tissue samples were homogenized in lysis buffer (Tris HCl pH 7.4 50 mM, NaCl 150 mM, EDTA 5 mM and 1 X-Triton X-100) containing phosphatase and protease inhibitors (Cocktail II, Sigma-Aldrich) to obtain total protein homogenates. For subcellular fractionation, 150 μL of buffer A (10 mM HEPES pH 7.9, 10 mM KCl, 0.1 mM EDTA pH 8, 0.1 mM EGTA pH 8, 1 mM DTT, 1 mM PMSF, protease inhibitors) were added to each sample and incubated on ice for 15 min. After this time, the samples were homogenized with a tissue homogenizer, 12.5 μL Igepal 1% were added, and mixed for 15 s. Following 30 s of full-speed centrifugation at 4°C, supernatants were collected (cytoplasmic fraction); 80 μL of buffer C (20 mM HEPES pH 7.9, 0,4M NaCl, 1 mM EDTA pH 8, 0.1 mM EGTA pH 8, 20% Glycerol 1 mM DTT, 1 mM PMSF, protease inhibitors) were added to each pellet and incubated under agitation at 4°C for 15 min. Subsequently, samples were centrifuged for 10 min at full speed at 4°C. Supernatants were collected (nuclear fraction) and 40 μL of buffer A+HCl (buffer A with 0.2 N HCl) were added to the pellet. After a 30-min incubation on ice, samples were centrifuged, again at full speed, at 4°C for 10 min and the supernatants were collected (the histone fraction). Aliquots of 15 μg of hippocampal protein extraction per sample were used. Protein samples were separated by Sodium dodecyl sulphate-polyacrylamide gel electrophoresis (SDS-PAGE) (8-14%) and transferred onto Polyvinylidene difluoride (PVDF) membranes (Millipore). Afterwards, membranes were blocked in 5% non-fat milk in Tris-buffered saline (TBS) solution containing 0.1% Tween 20 TBS (TBS-T) for 1 h at room temperature, followed by overnight incubation at 4°C with the primary antibodies listed in (Table S1). Then, the membranes were washed and incubated with secondary antibodies listed in (Table S1) for 1 h at room temperature. Immunoreactive proteins were viewed with the chemiluminescence-based ChemiLucent^™^ detection kit, following the manufacturer’s protocol (ECL Kit, Millipore), and digital images were acquired using ChemiDoc XRS+System (BioRad). Semi-quantitative analyses were done using ImageLab software (BioRad) and results were expressed in Arbitrary Units (AU), considering control protein levels as 100%. Protein loading was routinely monitored by immunodetection of Glyceraldehyde-3-phosphate dehydrogenase (GAPDH), β-tubulin or TATA-Binding protein (TBP).

### RNA extraction and gene expression determination by q-PCR

Total RNA isolation was carried out using TRIsure™ reagent according to the manufacturer’s instructions (Bioline Reagent, UK). The yield, purity, and quality of RNA were determined spectrophotometrically with a NanoDrop™ ND-1000 (Thermo Scientific) apparatus and an Agilent 2100B Bioanalyzer (Agilent Technologies). RNAs with 260/280 ratios and RIN higher than 1.9 and 7.5, respectively, were selected. Reverse Transcription-Polymerase Chain Reaction (RT-PCR) was performed as follows: 2 μg of messenger RNA (mRNA) was reverse-transcribed using the High Capacity cDNA Reverse Transcription Kit (Applied Biosystems). Real-time quantitative PCR (qPCR) was used to quantify mRNA expression of oxidative stress and inflammatory genes listed in (Table S2). SYBR^®^ Green realtime PCR was performed in a Step One Plus Detection System (Applied-Biosystems) employing SYBR^®^ Green PCR Master Mix (Applied-Biosystems). Each reaction mixture contained 6.75 μL of complementary DNA (cDNA) (which concentration was 2 μg), 0.75 μL of each primer (which concentration was 100 nM), and 6.75 μL of SYBR^®^ Green PCR Master Mix (2X).

Data was analysed utilizing the comparative Cycle threshold (Ct) method (ΔΔCt), where the housekeeping gene level was used to normalize differences in sample loading and preparation. Normalization of expression levels was performed with β-actin for SYBR^®^ Green-based real-time PCR results. Each sample was analysed in duplicate, and the results represent the n-fold difference of the transcript levels among different groups.

### Global DNA Methylation and Hydroxymethylation Determination

Isolation of genomic DNA was conducted using the FitAmp^™^ Blood and Cultured Cell DNA Extraction Kit (EpiGentek, Farmingdale, NY, USA) according to the manufacturer’s instructions. Following this, Methylflash Methylated DNA Quantification Kit (Epigentek, Farmingdale, NY, USA) and MethylFlash HydroxyMethylated DNA Quantification Kit were used in order to detect methylated and hydroxymethylated DNA. Briefly, these kits are based on specific antibody detection of 5-mC and 5-hmC residues, which trigger an ELISA-like reaction that allows colorimetric quantification at 450 nm.

### Oxidative Stress Determination

Hydrogen peroxide was measured in hippocampus protein homogenates as an indicator of oxidative stress, and it was quantified using the Hydrogen Peroxide Assay Kit (Sigma-Aldrich, St. Louis, MI) according to the manufacturer’s instructions.

### Data analysis

Data analysis was conducted using GraphPad Prism ver. 7 statistical software. Data are expressed as the mean ± standard error of the mean (SEM) of at least 6 samples per group. Diet and treatment effects were assessed by the Two-Way ANOVA analysis of variance, followed by Tukey post-hoc analysis or two-tail Student’s t-test when it was necessary. Statistical significance was considered when *p*-values were <0.05. The statistical outliers were determined with Grubbs’ test and subsequently removed from the analysis.

## 4. Results

### TAPS Assay

In TAPS assay the RL-118 target engagement was determined. The area of the peak relative to the number of cells was higher in cells that expressed the 11b-HSD1 enzyme (11β-HSD1 positive) compared to those cells that did not express the enzyme but were transfected with this gene (11β-HSD1 negative) (Fig. 1). In addition, differences between cells that expressed 11β-HSD1 positively and the Crimson positive cells were observed, indicating that the RL-118 drug is selective for its target.

**Figure 1.**
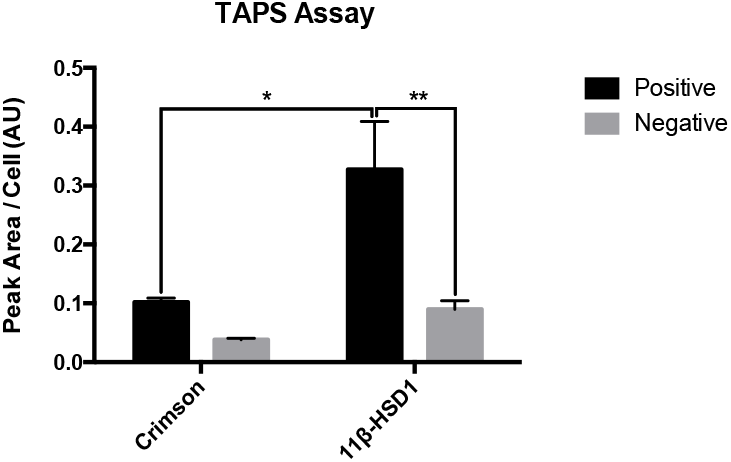
TAPS Assay results showing peak area per cell. Values are the mean ± Standard error of the mean (SEM) (n=2). *p<0.05; **p<0.01.

### CMS modulated epigenetic marks promoting an increased DNA methylation but reduced histone acetylation, reversed by 11β-HSD1 inhibition

Regarding DNA methylation, it can be observed that CMS reduced the overall DNA methylation and 5-hydroxy methylation compared to control group, although 11β-HSD1 inhibition increased those marks in both treated groups (Figs. 2 A-B). In accordance to the above-mentioned, DNA-methyltransferase 1 (Dnmt1) and ten-eleven translocase 2 (Tet2) gene expression were lower in CMS treated groups, while increased in control 11β-HSD1i treated group (Figs. 2 C-D).

**Figure 2.**
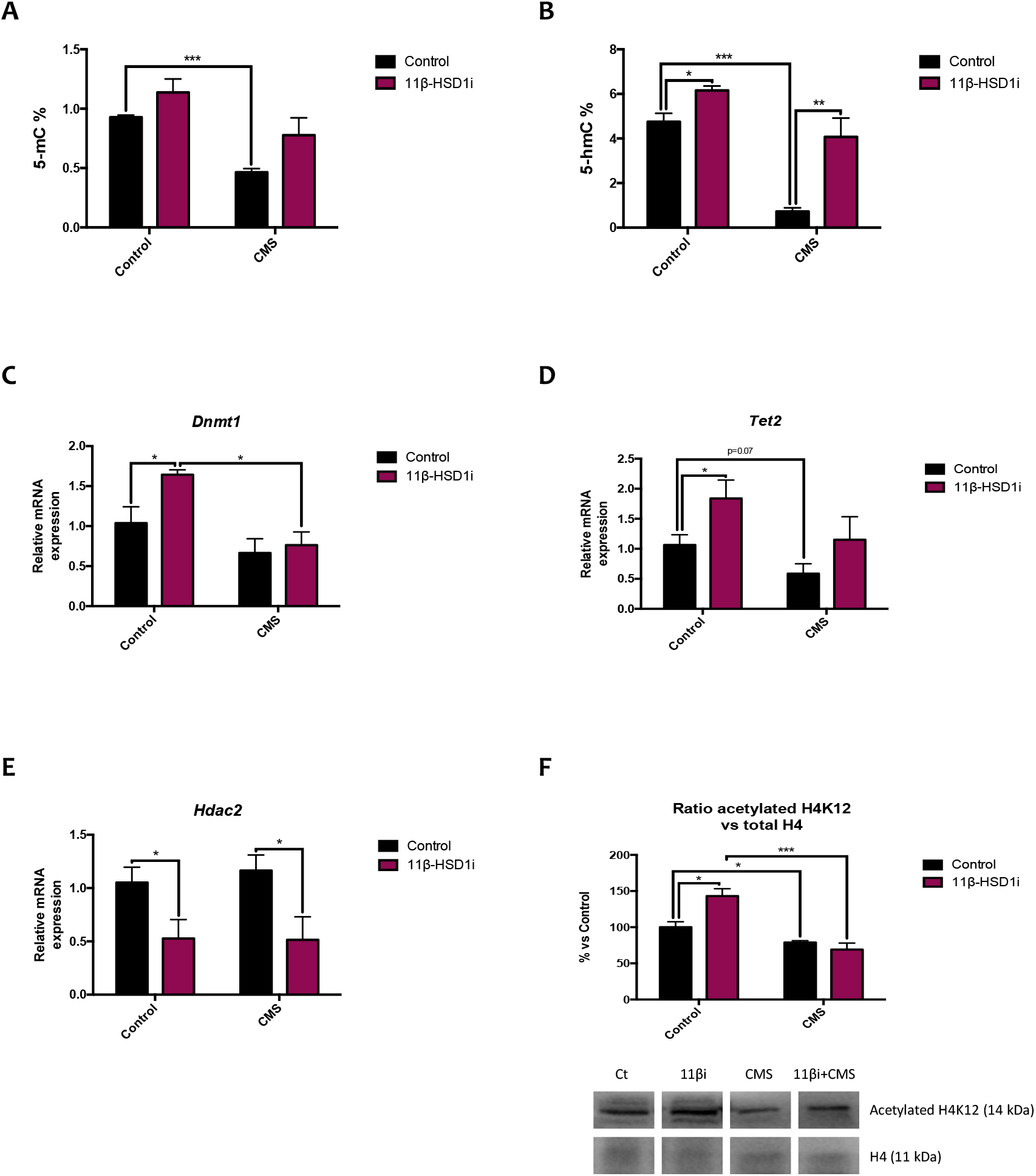
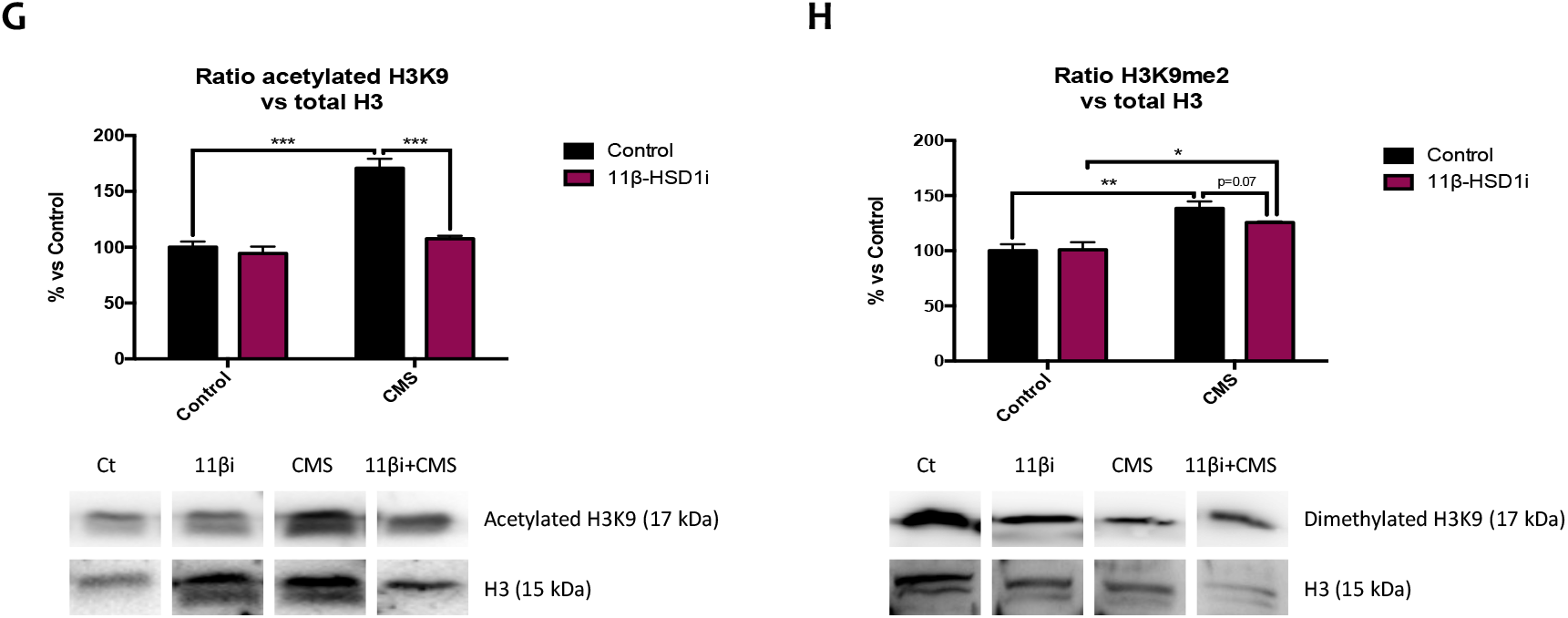
Representative results from epigenetic marks. Global 5-methylated cytosine (A) and 5-hydroxymethylated cytosine levels (B). Relative gene expression of Dnmt1 (C), Tet2 (D) and Hdac2 (E). Representative Western Blot for the ratio of Lys12 acetylated H4 protein levels and quantification (F), the ratio of Lys9 acetylated H3 protein levels and quantification (G) and the ratio of H3K9me2 protein levels and quantification (H). Gene expression levels were determined by real-time PCR. Western Blot values in bar graphs are adjusted to 100% for protein levels of SAMP8 Control (Control). Values are mean ± Standard error of the mean (SEM) (n = 6 for each group). *p<0.05; **p<0.01; ***p<0.001.

As far as histone epigenetic modifications are concerned, histone deacetylase 2 (Hdac2) gene expression was diminished by RL-118 treatment in both groups treated, while CMS seemed to increase it compared to control animals (Fig. 2 E). Mice under CMS showed lower Lys12 acetylated histone 4 protein levels and, accordingly, H4K12 protein levels were higher in 11β-HSD1i control group (Fig. 2 F). Lys9 acetylated histone 3 and di-methylation protein levels were higher in CMS group and accounting 11β-HSD1 inhibition effects, mice that also received CMS treatment showed higher H3K9me2 protein levels than the control mice (Figs. 2 G-H).

### GC levels attenuation, and conversely CMS application, led to decreased overall cellular OS

CMS increased ROS concentration in both groups compared to control mice. However, RL-118 drug treatment contributed to decrease ROS levels (Fig. 3 A). ROS accumulation is regulated by antioxidant and pro-oxidant enzymes controlled by nuclear erythroid-related factor 2 (Nrf2), such as heme oxygenase 1 (Hmox1) and Aldehyde oxidase 1 (Aox1). Hereby, it was found decreased Nrf2 protein levels after 11β-HSD1 inhibition, as well as increased protein levels in CMS treated groups (Fig. 3 B). In accordance, Aox1 gene expression was diminished in RL-118 treated groups, although increased after CMS treatment (Fig. 3 C). The same gene expression pattern was observed in for inducible nitric oxide synthase (iNOS), which is an enzyme implied in the synthesis of pro-oxidant molecules (Fig. 3 D).

**Figure 3.**
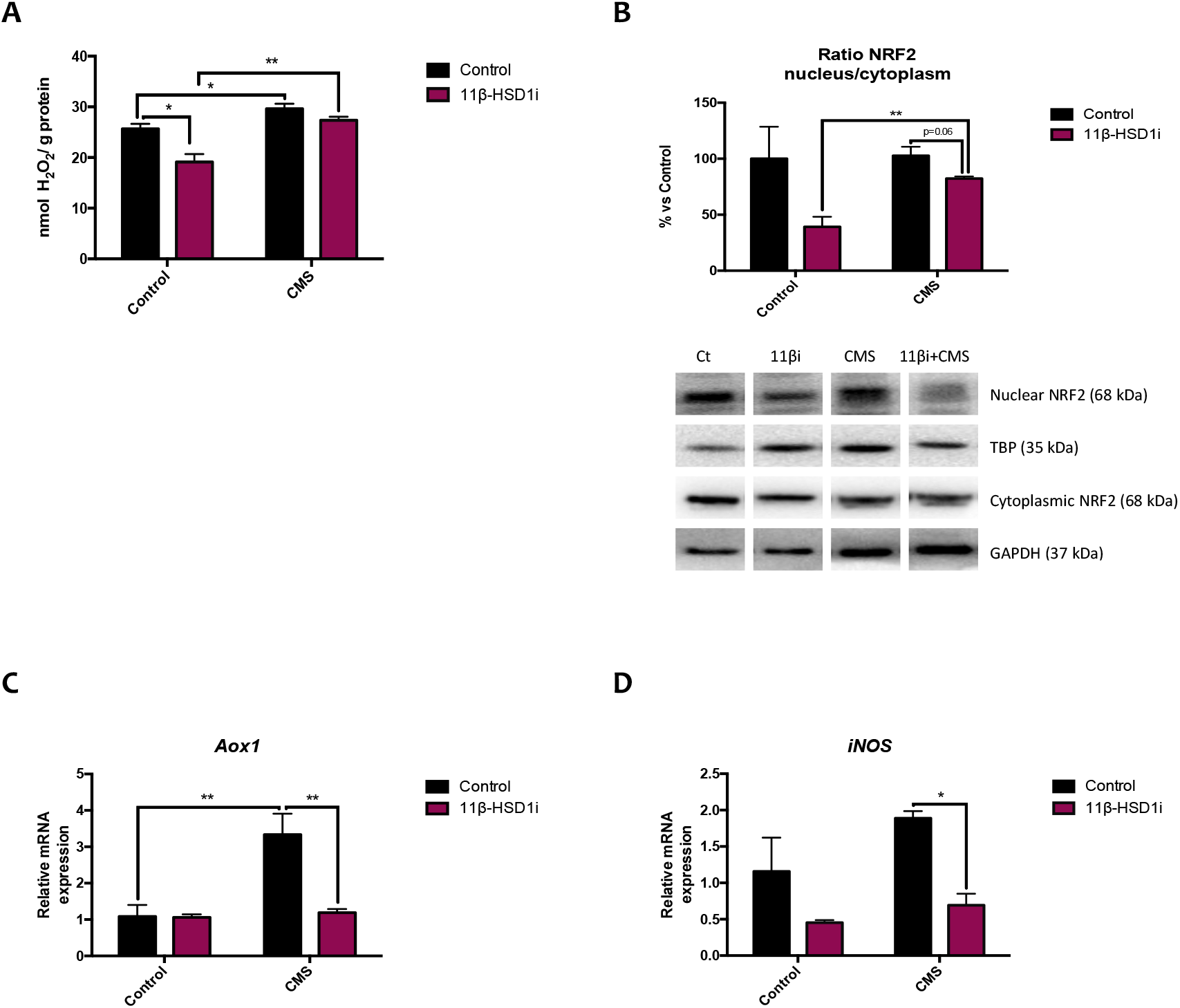
Representative results from pro-oxidant vs antioxidant mechanisms imbalance. Representative ROS accumulation measured as hydrogen peroxide concentration in homogenates of hippocampus tissue (A). Representative Western Blot for the ratio of nuclear/cytoplasmic NRF2 protein levels and quantification (B). Relative gene expression of Aox1 (C) and iNOS (D). Western Blot values in bar graphs are adjusted to 100% for protein levels of SAMP8 Control (Control). Gene expression levels were determined by real-time PCR. Values are mean ± Standard error of the mean (SEM) (n=6 for each group). *p<0.05; **p<0.01.

### Pro-inflammatory markers were reduced after 11β-HSD1 inhibition and CMS increased astrogliosis markers

Nuclear factor kappa-light-chain-enhancer of activated B cells (NF-κB) is a protein complex that mainly regulates cytokine production, thus inflammatory signalling and immune response to infection. Albeit no statistical differences were observed in NF-κB protein levels, there was a tendency to be decreased after RL-118 treatment in both treated groups (Fig. 4 A). However, there were differences in several cytokines gene expression controlled by this nuclear factor. Generally, CMS increased interleukin 1β (Il-1β), chemokine (C-X-C motif) ligand 2 (Cxcl-2) and tumour necrosis factor α (Tnf-α) gene expression, although not statistical differences; and 11β-HSD1 inhibition led to decreased cytokine gene expression (Figs. 4 B-D). Moreover, glial fibrillar acidic protein (Gfap) gene expression was evaluated and CMS increased its gene expression, which was reversed by 11β-HSD1 inhibition (Fig. 4 E).

**Figure 4.**
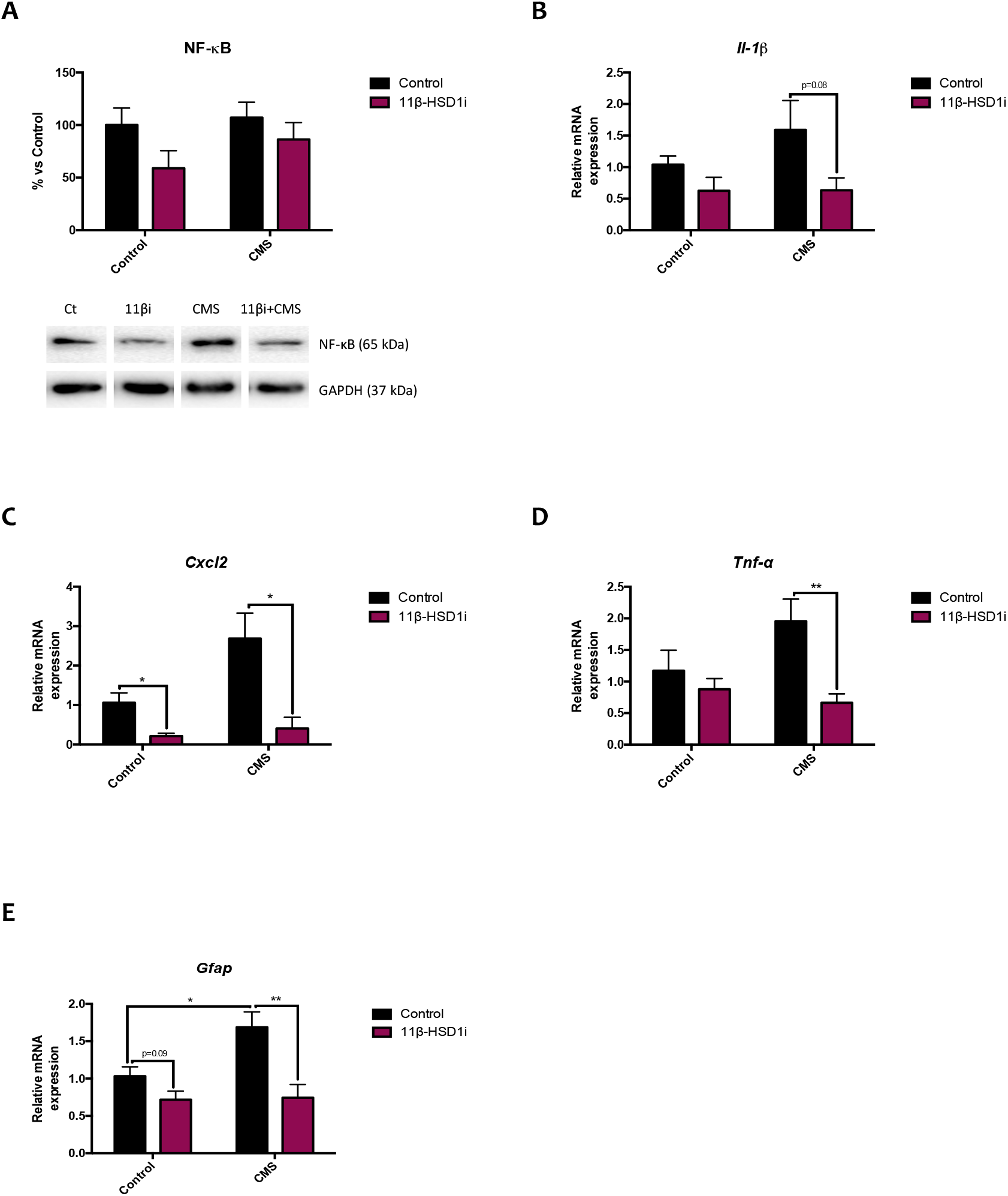
Representative results from inflammatory pathways. Representative Western Blot for NF-κB protein levels and quantification (A). Relative gene expression of Il-1β (B), Cxcl2 (C), Tnf-α (D) and Gfap (E). Western Blot values in bar graphs are adjusted to 100% for protein levels of SAMP8 Control (Control). Gene expression levels were determined by real-time PCR. Values are mean ± Standard error of the mean (SEM) (n=6 for each group). *p<0.05; **p<0.01.

### 11β-HSD1 inhibition and CMS promoted autophagy

Several autophagy markers were evaluated in the present work, such as Beclin1, rapamycin-sensitive TOR complex 1 (TORC1) and microtubule-associated protein 1A/1B light chain 3B (LC3B). Of note, Beclin1 and LC3B are activators of this cellular cleaning process, while TORC1 is associated with autophagy inhibition. After 11β-HSD1 inhibitor treatment, Beclin1 and LC3B protein levels were increased, while Ser151 phosphorylated TORC1 protein levels were decreased. Accordingly, mice under CMS showed higher protein levels of p-TORC1. However and strikingly, mice under CMS also presented higher protein levels of LC3B (Figs. 5 A-C).

**Figure 5.**
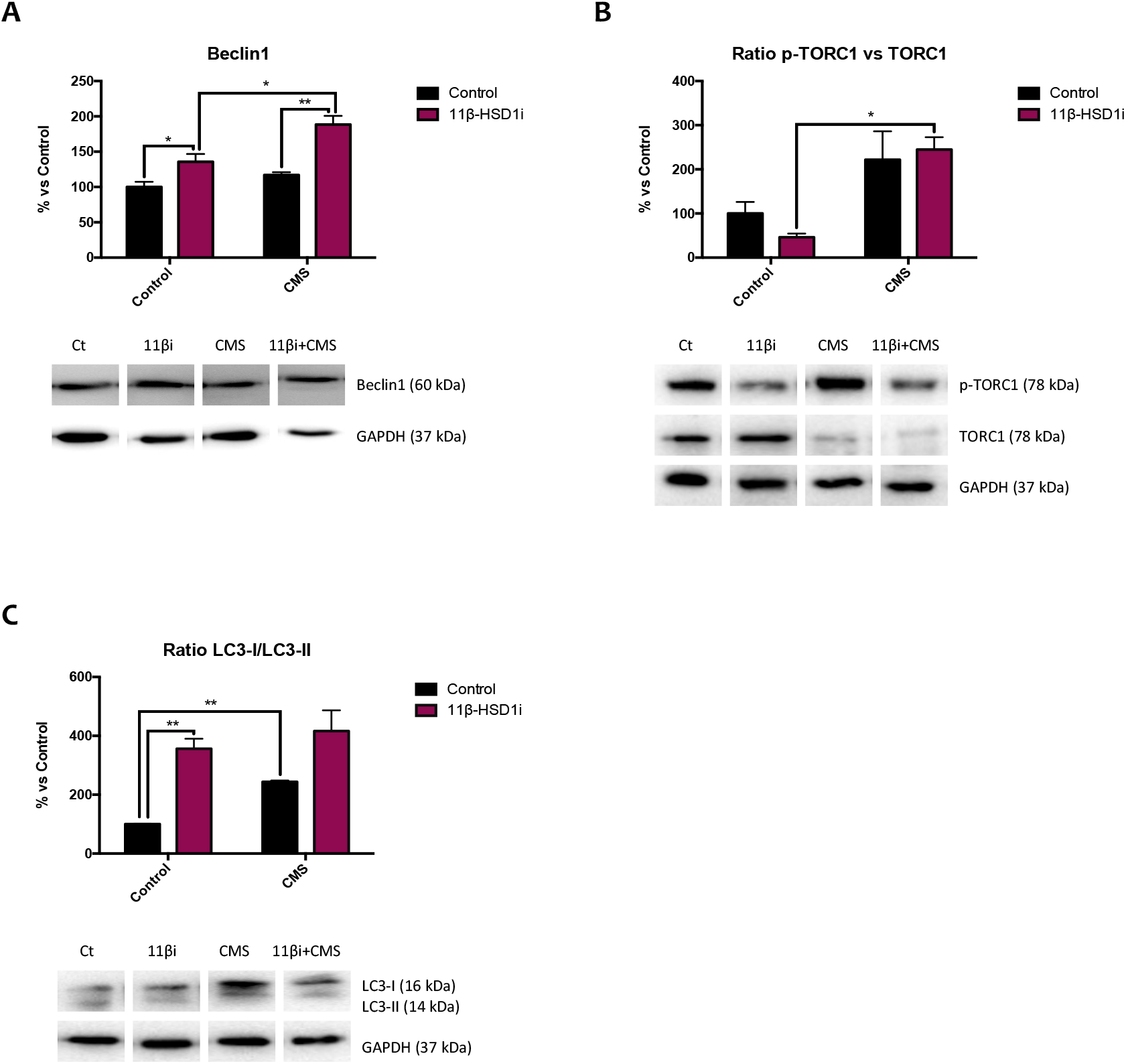
Representative results from autophagy process. Representative Western Blot for Beclin1 protein levels and quantification (A), the ratio of p-TORC1 protein levels and quantification (B) and the ratio of LC3 protein levels and quantification (C). Values in bar graphs are adjusted to 100% for protein levels of SAMP8 Control (Control). Values are mean ± Standard error of the mean (SEM) (n=4 for each group). *p<0.05; **p<0.01.

### 11β-HSD1 inhibition rescued mice from CMS pejorative effects on APP processing

Herein, a disintegrin and metalloproteinase domain protein 10 (Adam10) gene expression was increased in mice treated with RL-118 as well as decreased in CMS group (Fig. 6 A). Also, β-secretase 1 (Bace1) gene expression was decreased in both RL-118 treated groups and there was a trend towards increasing Bace1 gene expression after CMS exposure (Fig. 6 B). In line with these results, Aβ-precursor gene expression was increased in CMS group and reversed by 11β-HSD1 inhibition (Fig. 6 C). Finally, β-amyloid degradation was assessed by Neprilisin12 gene expression. It was found that RL-118 treatment increased its gene expression, while CMS decreased it (Fig. 6 D).

**Figure 6.**
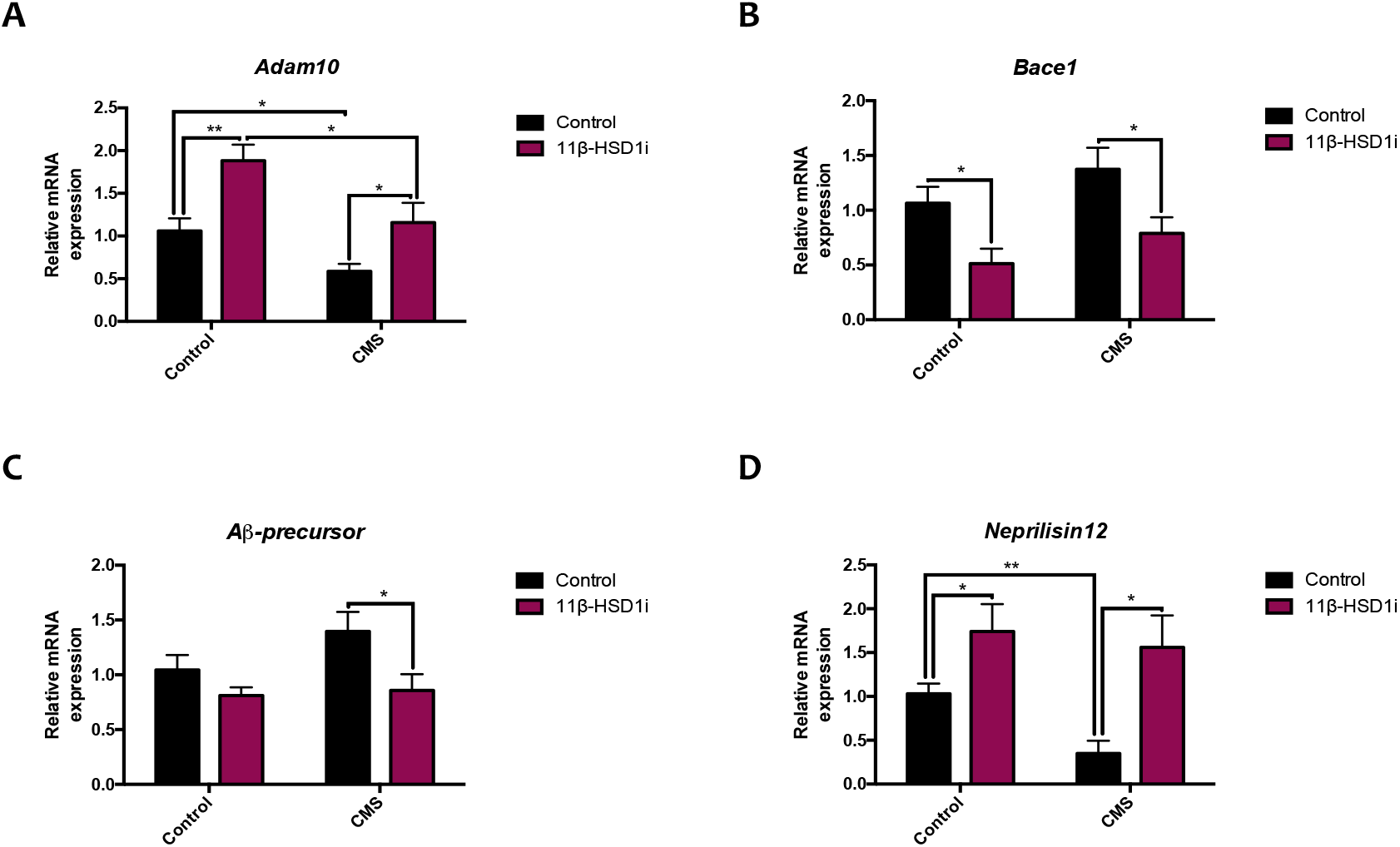
Representative results from APP processing pathways. Relative gene expression of Adam10 (A), Bace1 (B), Aβ-precursor (C) and Neprilisin12 (D). Gene expression levels were determined by real-time PCR. Values are mean ± Standard error of the mean (SEM) (n=6 for each group). *p<0.05; **p<0.01.

### Changes in synaptic plasticity were produced after GC attenuation

To address whether GC attenuation through 11β-HSD1 inhibition modulated synaptic plasticity, we evaluated different proteins involved in those mechanisms. cAMP response element-binding (Creb) gene expression and protein levels were increased in RL-118 treated mice and decreased after CMS exposure. As well as Creb’s downstream gene, brain-derived neurotrophic factor (Bdnf) (Figs. 7 A-C). Accordingly, synaptic modulators such as postsynaptic density protein 95 (PSD95), synaptophysin and synaptosomal nerve-associated protein 25 (SNAP25) protein levels were increased after 11β-HSD1 inhibition treatment, although not decreased after CMS (Figs. 7 E-H).

**Figure 7.**
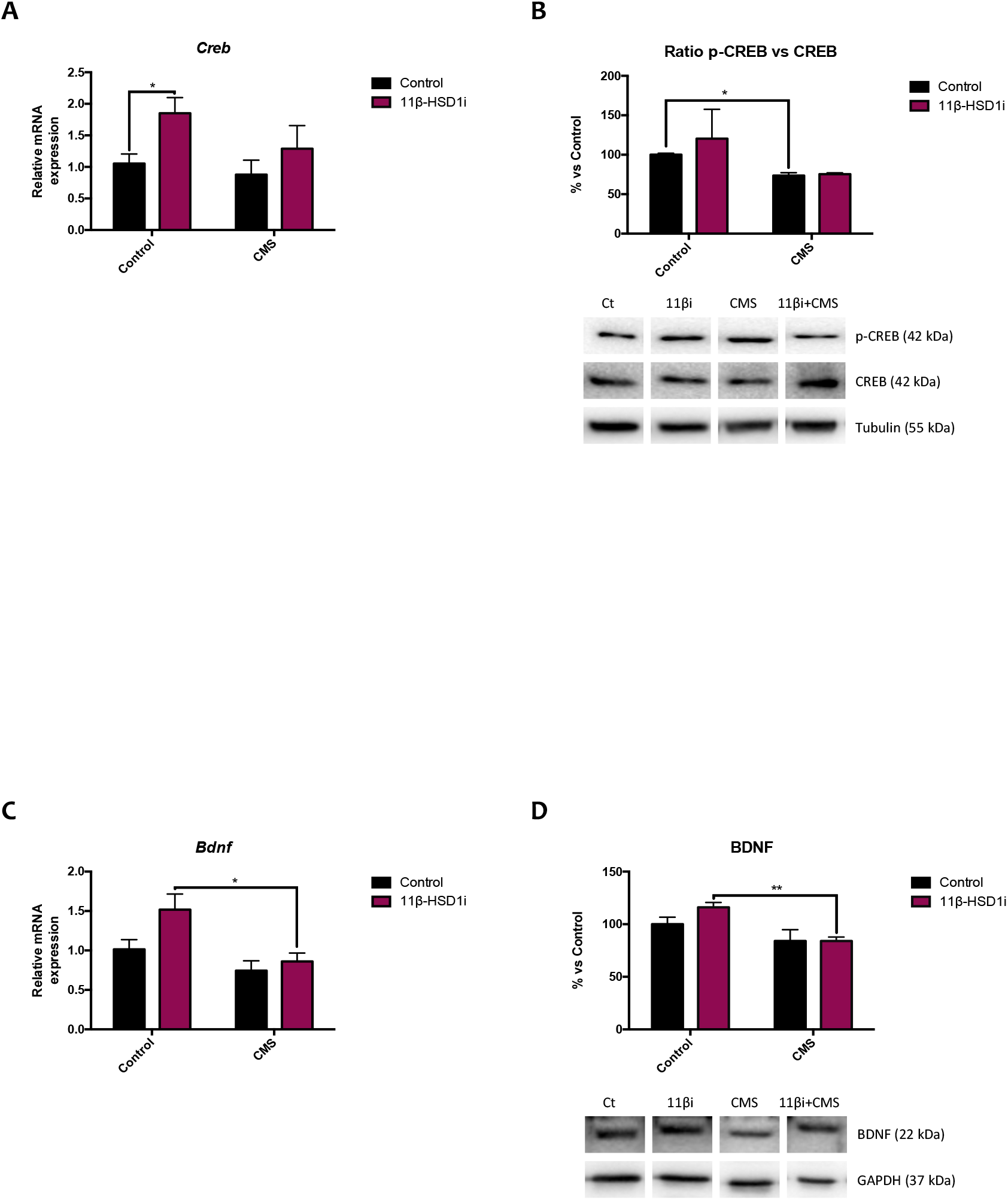
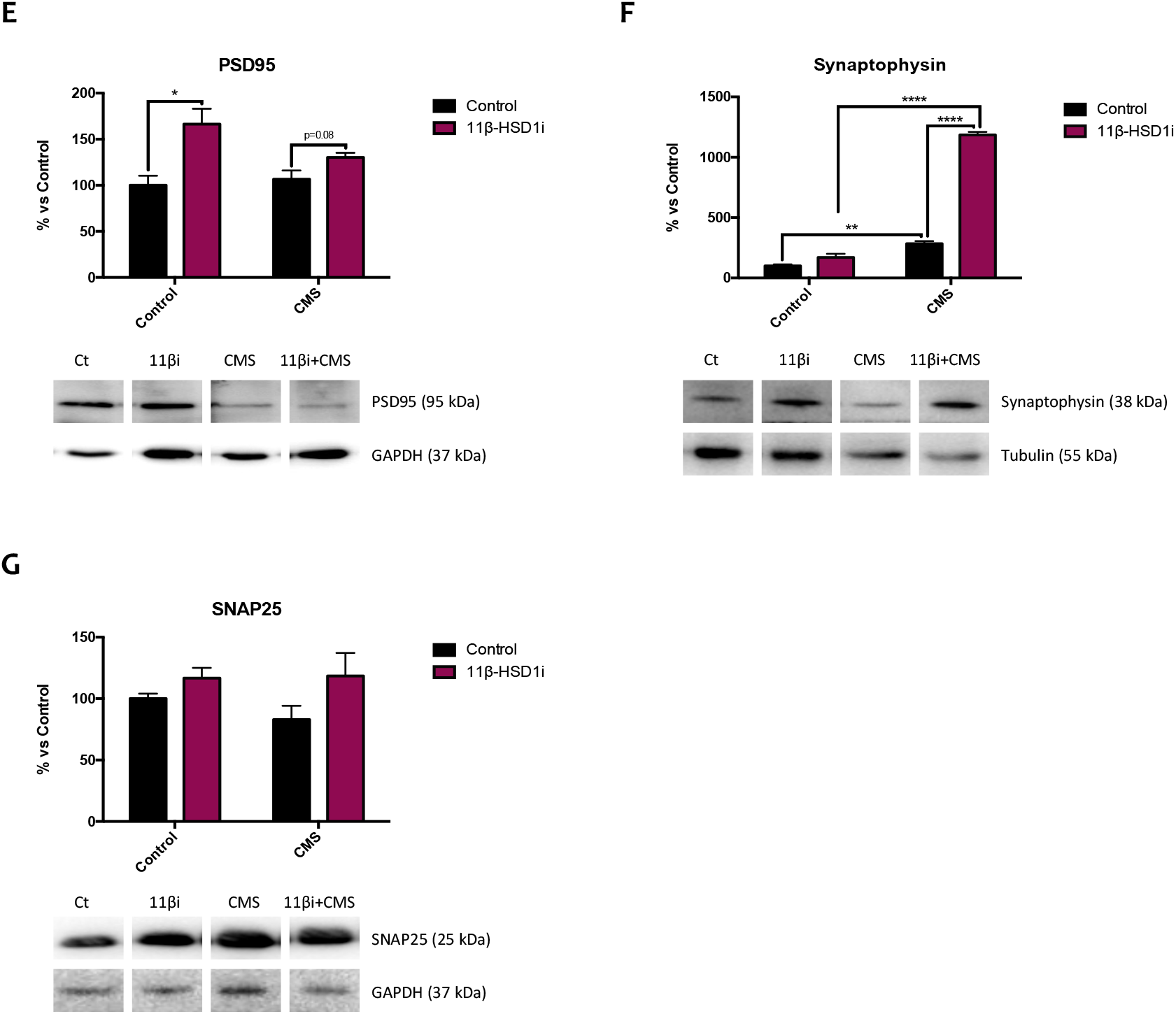
Representative results from neuroplasticity modulators. Relative gene expression of Creb (A). Representative Western Blot for the ratio of p-CREB protein levels and quantification (B). Relative gene expression of Bdnf (C). Representative Western Blot for BDNF protein levels and quantification (D), PSD95 protein levels and quantification (E), Synaptophysin protein levels and quantification (F) and SNAP25 protein levels and quantification (G). Gene expression levels were determined by real-time PCR. Western Blot values in bar graphs are adjusted to 100% for protein levels of SAMP8 Control (Control). Values are mean ± Standard error of the mean (SEM) (n=6 for each group). *p<0.05; **p<0.01; ***p<0.001; ****p<0.0001.

### 11β-HSD1 inhibition reduced risk-taking behaviour while increased memory and learning abilities and CMS vice verse

As mentioned, stress and GC influence behaviour and accelerate brain aging. CMS paradigm used did not change any parameters studied. However, 11β-HSD1 inhibition by RL-118 produced a tendency to increase horizontal but not vertical locomotor activity, while increased the time spent in the opened arms and decreased the time in the closed arms in the CMS mice, without affecting control groups (Figs. 8 A-D).

**Figure 8.**
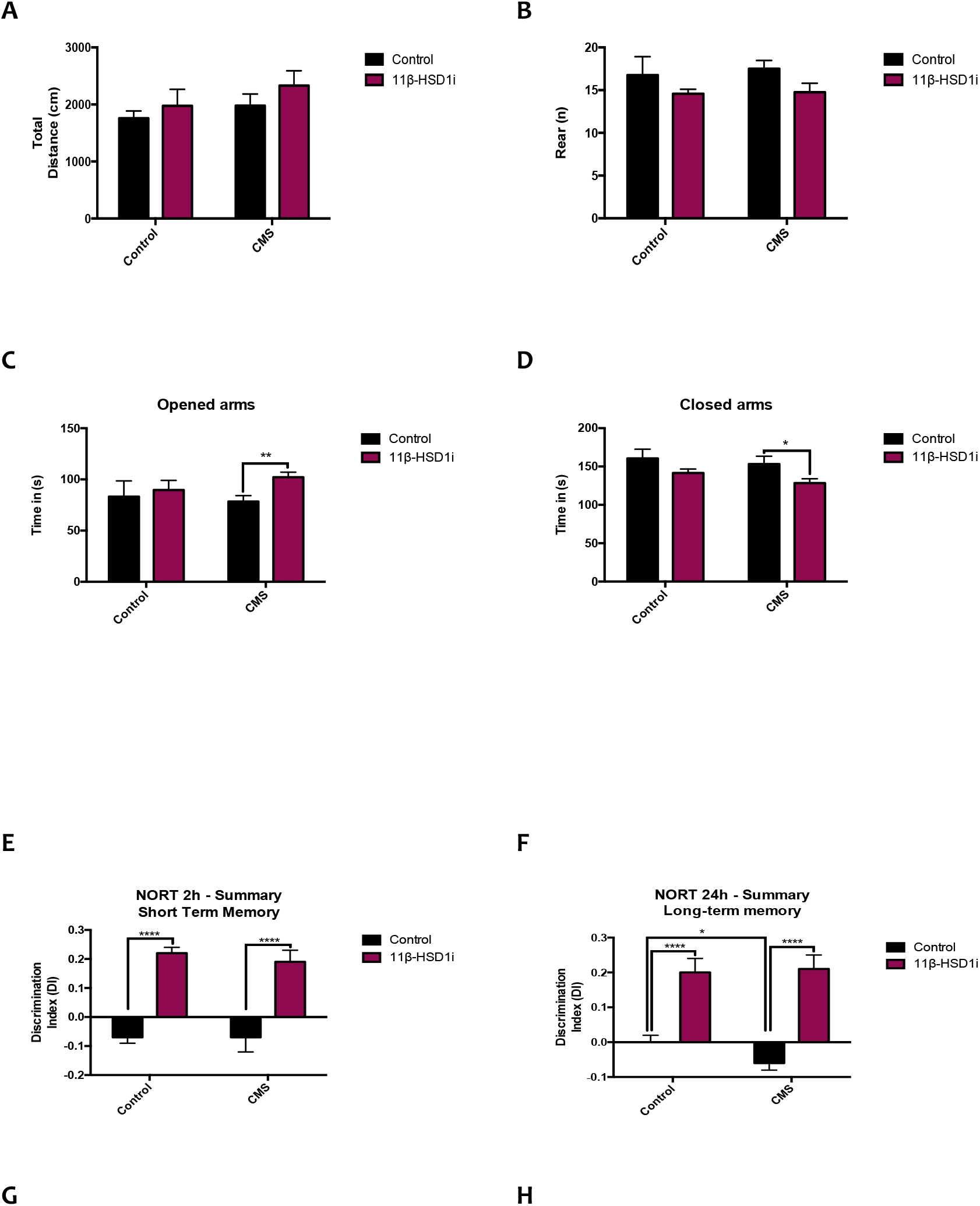
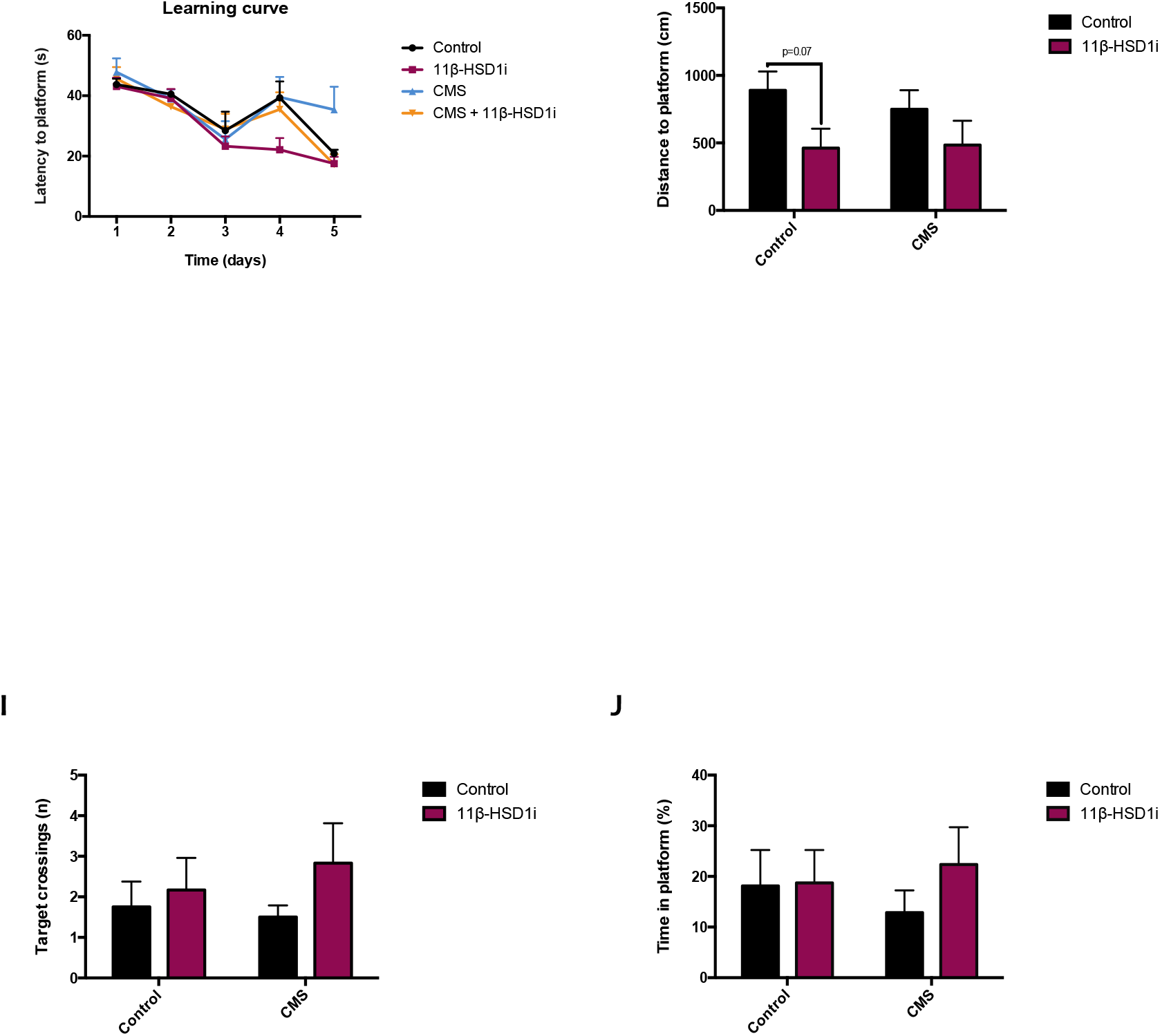
Behavioural tests results from EPM, NORT and MWM respectively. Total Distance (A), number of Rearing (B), Time in the Opened arms (C), Time in the Closed arms (D), Summary of DI from 2 and 24 hours after familiarization phase (E-F), MWM learning curve (G), Distance to reach the platform (H), number of Entries (I) and Time in the platform (J). Values are mean ± Standard error of the mean (SEM) (n=12 for each group). *p<0.05; **p<0.01; ***p<0.001; ****p<0.0001.

Referring to memory, we evaluated the recognition and spatial memory as well as learning abilities, through NORT and MWM. On one hand, CMS did not induce significant changes in the DI values in reference to control mice in NORT, although recognition memory was slightly reduced by CMS only after 24 hours familiarization (Figs. 8 E-F). Meanwhile, 11β-HSD1 inhibitor treatment increased recognition memory not only at short-term, but also at long-term. On the other hand, MWM learning curve demonstrates that all mice learned where the platform was during the training days, as the latency to the platform is lower than the first day although some groups performed better than others (Fig. 8 G). However, both groups treated with RL-118 showed better learning abilities in reference to CMS group, as the learning curve slope is higher in those groups. Overall, CMS treatment applied seemed to have little effect on the tests conducted, as there was a trend towards diminish all the parameters evaluated but not statistical differences were detected. By contrast, RL-118 treatment reduced the distance travelled to reach the platform, albeit there were only statistical significant differences between control groups (Fig. 8 H). In line with these results, 11β-HSD1 inhibition tended to increase target crossings as well as the time spent in the platform zone in both treated groups compared to their control littermates (Figs. 8 I-J).

## 5. Discussion

Taking into consideration the convergence of aging, stress and neurodegenerative diseases, such as AD, there is impaired GC signalling. Therefore, the study of GC response to chronic moderate stressful situations, as account in the daily life, results of huge interest in order to design pharmacological strategies to prevent neurodegeneration.

Several pieces of evidence demonstrate a strong association between prolonged exposure to GC excess and diminished cognitive abilities. It may be due to alterations in hippocampal electrophysiology, structure and function, as well as deleterious effects on neurotransmission, metabolism, cell division and death. GC secretion responds to a feed-forward signalling involving the hypothalamic-pituitary-adrenal (HPA) axis conforming the stress response. As far as stress is concerned, stressful stimuli increase GC secretion and release in order to confer energy to cope with the stress. Mounting evidence assert that chronic stress might lead to deleterious effects on brain, while acute stress may enhance memory (Sandi, 2013; Weger & Sandi, 2018). Accordingly, it has been demonstrated that chronic mild stress (CMS) paradigm in mice reduces cognitive abilities and increases anxiety-like behaviour (Puigoriol-Illamola et al., 2020a; Wang et al., 2016).

Previously, we affirmed that 11β-HSD1 enzyme regulates the disposure of active GC (Leiva et al., 2017; Puigoriol-Illamola et al., 2018). Not only that, but also 11β-HSD1 inhibition has been linked to enhance cognitive abilities and AD hallmarks (Mohler et al., 2011; Puigoriol-Illamola et al., 2018, 2020b). In line with these results, it has been described that 11β-HSD1 expression in mouse hippocampus and parietal cortex increases with aging and that its overexpression accelerated age-related cognitive decline (Holmes et al., 2010). By contrast, 11β-HSD1 knockout mice resist age-dependent cognitive loss (Yau et al., 2007, 2015). In our hands, we demonstrated that RL-118, a brain penetrant drug, treatment promoted autophagy flux, as well as ER stress activation in order to restore the deleterious effects exerted by prolonged exposure to GC (Leiva et al., 2017; Puigoriol-Illamola et al., 2018, 2020b). In conclusion, it is suggested that it could be a feasible target to fight against cognitive decline in age-related pathologies. In fact, early clinical studies demonstrate that 11β-HSD1 inhibitor (UE2343) is well tolerated in AD patients (Webster et al., 2017) and currently is at phase II. Besides, many selective 11β-HSD1 inhibitors have reached clinical stages for metabolic diseases, for instance AZD8329 and BVT.2733 (Puigoriol-Illamola et al., 2020b).

Up to now, studies demonstrating that RL-118 is able to reach the brain, bind the 11β-HSD1 enzyme and inhibit it, leading to therapeutic effects have not been addressed. In consequence, in the present work, the drug-target binding (Target engagement) of RL-118 and 11β-HSD1 was determined through a novel methodology consisting in FACS (fluorescence-activated cell sorting) coupled to mass spectrometry (MS) analysis (Wilson et al., 2017). Effectively, target engagement results showed that RL-118 drug binds to its target and also in a selectively way, since it did not bind to other proteins expressed. In consequence, we have determined that actually the molecular and behavioural effects observed after RL-118 administration are due to its binding to the 11β-HSD1 enzyme.

Stress response may have a genetic and epigenetic origin, reflected in the efficiency of the GC receptor (GR)-mediated GC negative feedback in the brain and/or the pituitary gland that causes HPA axis hyperactivity (Fink et al., 2017; Harman & Martin, 2019; Sotiropoulous et al., 2008). In an attempt to address whether and which changes produced the chronic presence of stressors we decided to evaluate two main epigenetic marks: DNA methylation and histone acetylation.

Regarding DNA methylation, it is implied in controlling neuronal gene expression and neural development. Dysregulation of this process is linked to a wide range of neuronal disorders, including AD onset and progression. However, the relationship between AD and altered 5-mC levels is not known (Fetahu et al., 2019; Puigoriol-Illamola et al., 2020a). TET family can further oxidize 5-mC to 5-hmC. Generally, 5-mC is associated with gene silencing, while 5-hmC with up-regulation of gene expression (Fetahu et al., 2019; Sherwani & Khan, 2015). Herein CMS reduced those epigenetic marks and accordingly to Puigoriol-Illamola et al. (2020a), attenuating GC levels led to increase 5-mC as well as 5-hmC percentages in both treated groups. Therefore, it may suggest that RL-118 is able to restore DNA methylation pattern although a detrimental stimulus such as CMS. Although these results seem to contradict what is established for most scientific articles, methylation of specific genes should be evaluated in order to know which translational activity is being repressed. Despite that, DNA hydroxymethylation, which is linked to promote gene expression, was increased in RL-118 treated mice. Overall, those results suggest that further evaluation of gene-promoter specific methylation are required to elucidate the clear epigenetic mechanisms to explain the neuroprotective effects observed in the cognitive tests performed. Likewise, the enzymes responsible for those processes as Dnmt1 and Tet2 gene expression were increased in both groups treated with 11β-HSD1 inhibitor, whereas it decreased in the CMS group.

DNA methylation and histone modifications must act coordinately (Zhao et al., 2016). Generally histone acetylation is related to favouring gene transcription through removing histone positive charge and thereby transforming condensed chromatin (heterochromatin) into a relaxed structure (euchromatin) (Turner, 2000; Watson et al., 2014). 11β-HSD1 inhibitor treatment altered Hdac2 gene expression, promoting a reduction in both groups. In line with this, acetylated H4 studied showed higher protein levels in RL-118 group treated, but not in mice under CMS too. In accordance to Puigoriol-Illamola et al. (2020a), SAMP8 mice under CMS treatment showed higher H3K9 protein levels in respect to control group. However, it could be reversed after concomitant RL-118 treatment. In turn, the same profile was observed in H3K9me2 protein levels evaluated, suggesting that although the increase in histone acetylation as it is also methylated, it finally results in transcriptional repression or a more compacted chromatin state in CMS groups compared to control mice.

Of note, although RL-118 is not a direct epigenetic modulator drug, it modifies the epigenetic landscape. Therefore, our results point out to the paramount importance of studying different drugs at epigenetic level in order to possess a deep knowledge of the pharmacological effects exerted by the drug.

Additional data (Kadmiel & Cidlowski, 2013; Picard et al., 2018; Puigoriol-Illamola et al., 2020a) report that a stressful environment affects oxidative balance. In normal conditions, pro-oxidant molecules and antioxidant defence mechanisms are balanced; however, under the presence of chronic stressors as well as along aging and in several disorders, there is a decrease in the capacity of the antioxidant enzymes allowing ROS accumulation and eventually causing cellular damage and finally dysfunction of the system (Griñán-Ferré et al, 2018). In concordance, in the present work SAMP8 mice under CMS increased ROS accumulation and on the contrary, those animals treated with RL-118 diminished it. OS is frequently invoked as a potential factor in progression of AD, yet whether it is a cause or consequence of the pathology is still debated (Bonet-Costa et al., 2016). Among antioxidant defence mechanisms, NRF2 pathway has been defined as a key indicator and modulator of OS in neurodegeneration (Griñán-Ferré et al., 2018; Johnson et al., 2008; Maeda et al., 2012), as it regulates the gene expression of different antioxidant enzymes. While NRF2 is a protector against oxidative and electrophilic tissue injury, persistent activation of NRF2 signalling may also contribute to disease pathophysiology (Li et al., 2019). Interestingly, NRF2 protein levels were decreased after RL-118 treatment, suggesting that attenuating GC levels induce lower oxidative damage. However, CMS stimulated the increase in Aox1 gene expression, which was reduced after concomitant RL-118 treatment, indicating that stressful environment increases pro-oxidant mechanisms while 11β-HSD1 inhibition is able to restore them. Likewise, other OS modulators evaluated as iNOS agreed with the increase in CMS group compared to control group and decreased protein levels of gene expression after RL-118 treatment. Moreover, iNOS gene expression has been reported to be regulated by inflammatory signalling, particularly through NF-κB (Arias-Salvatierra et al., 2011). Overall, these results suggest that 11β-HSD1 inhibition promoted antioxidant mechanisms to cope with OS in a mice model of aging, even under CMS conditions.

GCs effects include inflammatory signalling regulation. While normal GC activity involves immune system suppression, GC excess deploys the contrary effect. Despite differences did not reach statistical significance, 11β-HSD1 inhibition reduced NF-κB protein levels. It appears to be a pivotal mediator of inflammatory responses as it induces the expression of various pro-inflammatory genes, including those encoding cytokines and chemokines, and participates in inflammasome regulation (Liu et al., 2017). It triggers a feed-forward cycle in which increases cytokine production and this in turn result in resistance to the GC immunosuppression, leading to further increases in cytokine release and, therefore, activation of the HPA axis (Irwin & Miller, 2007; Pace et al., 2007). Accordingly, we found decreased pro-inflammatory cytokine gene expression after 11β-HSD1 inhibitor treatment and increased after CMS appliance. Moreover, Il-10 and Il-6 gene expression results reinforce the same assertion (Data not shown). Pro-inflammatory cytokines like IFN-α, Il-1, IL-6 and TNF-α have been described to activate the HPA axis and potentiate GC resistance (Sotiropoulous et al., 2008). Additionally, they have been implicated in the genesis of AD, becoming part of the amyloid plaques or triggering amyloid-β (Aβ) production, among others. Dysregulation of inflammatory mediators and astrogliosis are major culprits in the development of chronic inflammation and immunosenescence process, as well as are related to cognitive decline and progression of neurodegenerative diseases (Chung et al., 2019).

Recent studies have indicated that prolonged OS may limit autophagy flux (Bonet-Costa et al., 2016). In addition, altered cellular loss of proteostasis is one of the nine hallmarks of aging postulated by López-Otín et al. (2013). Moreover, our group had previously demonstrated that attenuating GC excess by RL-118 promotes autophagy activation (Puigoriol-Illamola et al., 2018) and, by contrast, CMS treatment induced autophagy deterioration (Puigoriol-Illamola et al., 2020a). In accordance, autophagy flux was decreased in female mice under CMS but those treated with RL-118 showed higher protein levels of autophagy activators, like Beclin1 and LC3B, and lower protein levels of p-TORC1.

11β-HSD1 inhibition has been linked to an extensive number of disorders, among which there is AD. In particular, it has been associated with reduced Aβ neurotoxicity and Tau hyperphosphorylation (Dong & Csernansky, 2009; Ouanes & Popp, 2019), both hallmarks of AD. Analysing deeper this issue, amyloid precursor protein (APP) can be processed through two mechanisms: non-amyloidogenic and amyloidogenic pathways. The main indicator of the former is Adam10, whereas the later is Bace1 (Shen et al., 2018). In this study, Adam10 gene expression was diminished and Bace1 increased after CMS. Importantly, in RL-118 treated groups occurred the opposite, then favouring the slow down formation of Aβ. Moreover, the Aβ degrading enzyme, neprilisin12 gene expression increased after 11β-HSD1 inhibitor treatment but reduced after CMS.

As mentioned above, prolonged exposure to GCs in the brain induces several changes and ample experimental evidence state that repeated exposure to stressful conditions induce structural remodelling of neurons with synaptic loss as well as alterations in glial functions, which are frequently maladaptive (Vyas et al., 2016). In agreement, our results demonstrate a reduction in Creb and Bdnf gene expression and protein levels after CMS exposure, but conversely RL-118 treatment was not able to prevent it (Steffke et al., 2020). In a similar way, CMS reduced synaptic plasticity markers, like PSD95, Synaptophysin and SNAP25, and by contrast 11β-HSD1 inhibition increased them, in particular Synaptophysin. Thus, RL-118 treatment could prevent the loss of those neuroplasticity markers, in particular Synaptophysin (Mango et al., 2019).

The effects of GCs on cognition have been widely studied, stating that acute stress improves cognitive abilities, while chronic stress worsens memory and learning processes as well as accelerates brain aging (Ouanes & Popp, 2019; Sandi et al., 2013; Sotiropoulous et al., 2008). In fact, GC levels have been found to correlate with the severity of the cognitive impairment (Ouanes & Popp, 2019). Assessing EPM results, it is well known that vertical locomotor activity indicates anxiety-like behaviour and, although not statistical differences were observed, together with the total horizontal distance travelled, we could affirm that RL-118 drug treatment helped to reduce feeling of anxiety and desire to escape from an inhospitable environment, so that risk-taking behaviour becomes reduced. This is supported by the time spent in opened and closed arms, as it can be seen that after 4 weeks of 11β-HSD1 inhibition together to detrimental stimuli such as CMS, animals stood less time in the closed arms and, on the contrary, more time in the opened arms respect to CMS group. In accordance to Puigoriol-Illamola et al. (2020 a & b), mice under RL-118 treatment clearly showed improved recognition memory than control mice. However, the pejorative effect caused by CMS was only detected at long-term recognition memory evaluation. Regarding MWM results, it seemed that CMS treatment might not be strong enough to produce clearly changes in the tests performed. 11β-HSD1 inhibition tended to improve the mice performance, although not statistical differences were detected. However, both groups treated with RL-118 showed better learning abilities in reference to CMS group, as the learning curve slope is higher in those groups, indicative of a putative role for RL-118 as a neuroprotectant or cognitive enhancer of cognition (nootropic drugs).

## 6. Conclusions

In view of these results, target engagement between 11β-HSD1 enzyme and RL-118 drug was demonstrated and therefore, we can surely attribute all the beneficial effects observed in SAMP8 treated with RL-118 to 11β-HSD1 inhibition. Diversely, CMS declined cognitive and behavioural abilities, as well as synaptic plasticity, autophagy and antioxidant mechanisms, while modulated epigenetic markers and increased inflammatory signalling and Aβ formation and accumulation. However and most important, 11β-HSD1 inhibition through RL-118 turned up to restore the majority of these detrimental effects caused by CMS (Fig. 9).

**Figure 9.**
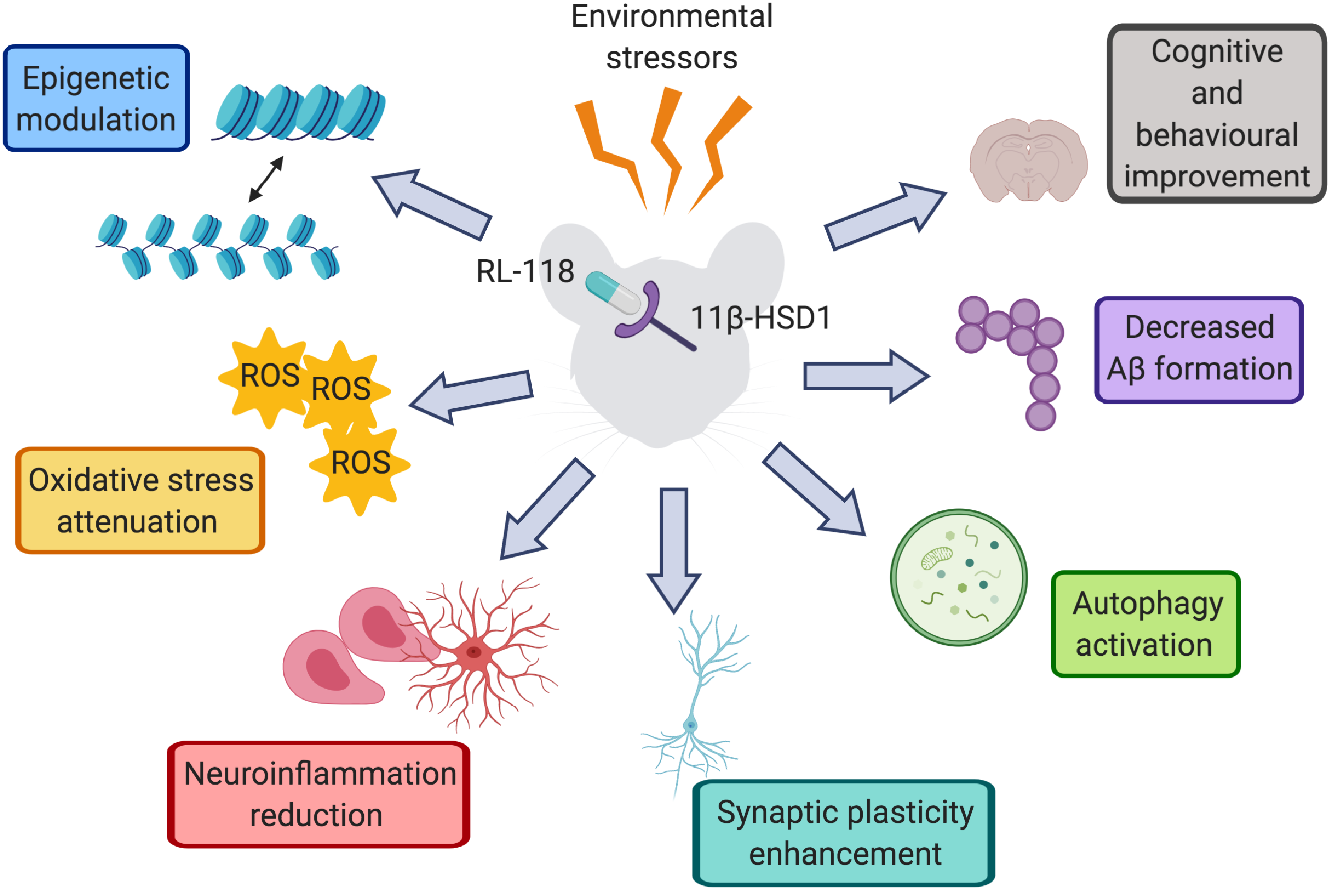
Representative scheme of the molecular pathways altered after RL-118 treatment in SAMP8 mice under chronic mild stress.

## 7. Acknowledgements

This study was supported by Ministerio de Economía y Competitividad of Spain and FEDER (SAF2016-77703) and 2017SGR106 (AGAUR, Catalonia). Financial support was provided for D.P.I. (FPU program).

