## Supplemental Material 1, Supplemental Table 1, Supplemental Table 2 for "RL-118 and 11β-HSD1 target engagement through TAPS assay: behaviour and molecular analysis"

### 3.3.9 Supplementary material

**Fig. S1 E2-Crimson-huHSD11B1 fusion protein.** Restriction sites for NheI and NotI are marked in pink, Kozak sequence in bold, E2-Crimson gene in red, a linker sequence in blue and HSD11B1 gene in black.

**NheI** **Kozak**
**5’GCTAGCGCCACCATGGCGAGCATG**GATAGCACTGAGAACGTCATCAAGCCCTTCATGCGCTTCAA
GGTGCACATGGAGGGCTCCGTGAACGGCCACGAGTTCGAGATCGAGGGCGTGGGCGAGGGCAAGC
CCTACGAGGGCACCCAGACCGCCAAGCTGCAAGTGACCAAGGGCGGCCCCCTGCCCTTCGCCTGGG
ACATCCTGTCCCCCCAGTTCTTCTACGGCTCCAAGGCGTACATCAAGCACCCCGCCGACATCCCCGAC
TACCTCAAGCAGTCCTTCCCCGAGGGCTTCAAGTGGGAGCGCGTGATGAACTTCGAGGACGGCGGC
GTGGTGACCGTGACCCAGGACTCCTCCCTGCAGGACGGCACCCTCATCTACCACGTGAAGTTCATCG
GCGTGAACTTCCCCTCCGACGGCCCCGTAATGCAGAAGAAGACTCTGGGCTGGGAGCCCTCCACTG
AGCGCAACTACCCCCGCGACGGCGTGCTGAAGGGCGAGAACCACATGGCGCTGAAGCTGAAGGGC
GGCGGCCACTACCTGTGTGAGTTCAAGTCCATCTACATGGCCAAGAAGCCCGTGAAGCTGCCCGGCT
ACCACTACGTGGACTACAAGCTCGACATCACCTCCCACAACGAGGACTACACCGTGGTGGAGCAGT
ACGAGCGCGCCGAGGCCCGCCACCACCTGTTCCAG**GATGACGATGACAAGATG**GCTTTTATGAAAA
AATATCTCCTCCCCATTCTGGGGCTCTTCATGGCCTACTACTACTATTCTGCAAACGAGGAATTCAGA
CCAGAGATGCTCCAAGGAAAGAAAGTGATTGTCACAGGGGCCAGCAAAGGGATCGGAAGAGAGAT
GGCTTATCATCTGGCGAAGATGGGAGCCCATGTGGTGGTGACAGCGAGGTCAAAAGAAACTCTACA
GAAGGTGGTATCCCACTGCCTGGAGCTTGGAGCAGCCTCAGCACACTACATTGCTGGCACCATGGA
AGACATGACCTTCGCAGAGCAATTTGTTGCCCAAGCAGGAAAGCTCATGGGAGGACTAGACATGCT
CATTCTCAACCACATCACCAACACTTCTTTGAATCTTTTTCATGATGATATTCACCATGTGCGCAAAA
GCATGGAAGTCAACTTCCTCAGTTACGTGGTCCTGACTGTAGCTGCCTTGCCCATGCTGAAGCAGAG
CAATGGAAGCATTGTTGTCGTCTCCTCTCTGGCTGGGAAAGTGGCTTATCCAATGGTTGCTGCCTATT
CTGCAAGCAAGTTTGCTTTGGATGGGTTCTTCTCCTCCATCAGAAAGGAATATTCAGTGTCCAGGGT
CAATGTATCAATCACTCTCTGTGTTCTTGGCCTCATAGACACAGAAACAGCCATGAAGGCAGTTTCT
GGGATAGTCCATATGCAAGCAGCTCCAAAGGAGGAATGTGCCCTGGAGATCATCAAAGGGGGAGC
TCTGCGCCAAGAAGAAGTGTATTATGACAGCTCACTCTGGACCACTCTTCTGATCAGAAATCCATGC
AGGAAGATCCTGGAATTTCTCTACTCAACGAGCTATAATATGGACAGATTCATAAACAAG**TAGGCG**
**GCCGC3’**
**Not1**

**Table S1. Antibodies used in Western Blot studies.**

| Antibody | Host | Source/Catalog | WB dilution |
|---|---|---|---|
| **Acetyl H4K12** | Sheep | R&D Systems/AF5215 | 1:1000 |
| **H4** | Rabbit | Cell Signaling/#2592 | 1:1000 |
| **Acetyl H3K9** | Rabbit | Millipore/06-599 | 1:1000 |
| **H3** | Rabbit | Cell Signaling/#9715 | 1:1000 |
| **H3K9me2** | Rabbit | Cell Signaling/#4658 | 1:1000 |
| **NF-κβ** | Rabbit | Cell Signaling/DE14E12 | 1:1000 |
| **NRF2** | Rabbit | Santa Cruz/sc-722 | 1:500 |
| **Beclin 1** | Rabbit | Abcam/ab62557 | 1:1000 |
| **p-TORC1 S151** | Rabbit | Cell Signaling/#3359 | 1:1000 |
| **TORC1** | Rabbit | Cell Signaling/#2501 | 1:1000 |

| | | | |
|---|---|---|---|
| **LC3B** | Rabbit | Cell Signaling/#2775 | 1:1000 |
| **p-CREB S133** | Rabbit | Cell Signaling/#9198 | 1:1000 |
| **CREB** | Rabbit | Cell Signaling/#4820 | 1:1000 |
| **BDNF** | Rabbit | Santa Cruz/N-20 | 1:500 |
| **PSD95** | Rabbit | Abcam/ab12093 | 1:2000 |
| **Synaptophysin** | Mouse | Millipore/MAB5258 | 1:1000 |
| **SNAP25** | Mouse | Santa Cruz/ SP-12 | 1:500 |
| **GAPDH** | Mouse | Millipore/MAB374 | 1:2000 |
| **β-Tubulin** | Mouse | Millipore/Clone AA2 | 1:2000 |
| **TBP** | Mouse | Abcam/ab818 | 1:2000 |
| **Goat-anti-mouse HRP conjugated** | | Biorad/170-5047 | 1:2000 |
| **Goat-anti-rabbit HRP conjugated** | | Biorad/170-6515 | 1:2000 |
| **Donkey-anti-goat HRP conjugated** | | Santa Cruz/sc-2020 | 1:2000 |

**Table S2. Primers and probes used in qPCR studies.**

**SYBR Green primers**

| Target | Product size (bp) | Forward primer (5'-3') | Reverse primer (5'-3') |
|---|---|---|---|
| ***Tet2*** | 113 | CCATCATGTTGTGGGACGGA | ATTCTGAGAACAGCGACGGT |
| ***Hdac2*** | 280 | CTATCCCGCTCTGTGCCCT | GAGGCTTCATGGGATGACCC |
| ***G9a*** | 94 | TTCCTTGTCTCCCCTCCCAG | CTATGAACTCTCTCGGCGGC |
| ***Aox1*** | 286 | CATAGGCGGCCAGGAACATT | TCCTCGTTCCAGAATGCAGC |
| ***iNOS*** | 101 | GGCAGCCTGTGAGACCTTTG | GAAGCGTTTCGGGATCTGAA |
| ***Il-1β*** | 179 | ACAGAATATCAACCAACAAGT TGATATTCTC | GATTCTTTCCTTTGAGGCCCA |
| ***Cxcl2*** | 100 | AGCCACACTTCAGCCTAGCG | TGTAGCCTGGTGGTTGGTGG |
| ***Tnf-α*** | 157 | TCGGGGTGATCGGTCCCCAA | TGGTTTGCTACGACGTGGGCT |
| ***Gfap*** | 125 | CCTTCTGACACGGATTTGGT | ACATCGAGATCGCCACCTAC |
| ***Adam10*** | 125 | GGGAAGAAATGCAAGCTGAA | CTGTACAGCAGGGTCCTTGAC |
| ***Bace1*** | 67 | ACAAGCCTTTCCGCCTCC | TCAGGCCACCATAATCCAGC |
| ***Aβ-precursor*** | 99 | TCGGGGTGATCGGTCCCCAA | GTCACGTTCACCCTCCCCAG |
| ***Neprilisin12*** | 196 | TTGGGAGACCTGGCGGAAAC | CATTCCTTGGACCCTCACCCC |
| ***Creb*** | 86 | GGCTGCTGCTGCCTGT | ACACACCGCGTCAAACTACA |
| ***Bdnf*** | 72 | TAGCTTGACAAGGCGAAGGG | TCTGGCAAAGATGAGCTCGG |
| ***β-actin*** | 190 | CAACGAGCGGTTCCGAT | GCCACAGGTTCCATACCCA |

**Taqman probes**

| Target | Product size (bp) | Reference |
|---|---|---|

| | | |
|---|---|---|
| ***Dnmt1*** | 58 | Mm01151063_m1 |
| ***Gapdh*** | 107 | Mm99999915_g1 |
